# Resident Tissue Macrophages Govern Intraocular Pressure Homeostasis

**DOI:** 10.1101/2025.01.25.634888

**Authors:** Katy C. Liu, Aleksander O. Grimsrud, Maria Fernanda Suarez, Darren Schuman, Michael L. De Ieso, Megan Kuhn, Ruth A. Kelly, Rose Mathew, Joan Kalnitsky, Matthias Mack, Florent Ginhoux, Violet Bupp-Chickering, Revathi Balasubramanian, Simon W. M. John, W. Daniel Stamer, Daniel R. Saban

## Abstract

Intraocular pressure is tightly regulated by the conventional outflow tissues, preventing ocular hypertension that leads to neurodegeneration of the optic nerve, or glaucoma. Although macrophages reside throughout the conventional outflow tract, their role in regulating intraocular pressure remains unknown. Using macrophage lineage tracing approaches, we uncovered a dual macrophage ontogeny with distinct spatial organizations across the mouse lifespan. Long-lived, resident tissue macrophages concentrated in the trabecular meshwork and Schlemm’s canal, whereas short-lived monocyte-derived macrophages, instead, were abundant around distal vessels. Specific depletion of resident tissue macrophages triggered elevated intraocular pressure and outflow resistance, linked to aberrant extracellular matrix turnover in the resistance-generating tissues of the trabecular meshwork. This dysregulated physiology and tissue remodeling were not observed when we depleted monocyte-derived macrophages. Results show ontogeny and tissue-specific macrophage function within the outflow tract, uncovering the integral homeostatic role of resident tissue macrophages in resistance-generating tissues whose dysfunction is responsible for glaucoma.

## INTRODUCTION

Glaucoma is the second leading cause of blindness worldwide ^1^. Despite effective treatments that work by lowering intraocular pressure (IOP) ^2–4^, some patients continue to lose vision due to inadequate IOP control ^5,6^. The significance of the immune system in IOP regulation has been widely recognized (i) due to the effect of specific ocular treatments on both immune cells and IOP ^7–10^, (ii) the abundance of macrophages in the tissues that regulate IOP ^8,11^, (iii) and genome-wide association studies showing significant changes in gene-sets involving the immune system in glaucoma patients ^12^. However, there has been no direct experimental evidence showing that the immune system contributes to IOP homeostasis, which is maintained within 1-2 mmHg in healthy people by continual adjustments in outflow resistance over a lifetime ^13–15^.

To lower IOP in glaucoma patients, current therapies exploit the molecular mechanisms that control both inflow and outflow of aqueous humor. First-line glaucoma treatment uses laser energy directed towards outflow tissues to reduce IOP ^16^, with concurrent monocyte recruitment ^8^. In addition, another first line anti-glaucoma treatment, topical prostaglandin F2α receptor agonists, lowers IOP ^17^ by altering extracellular matrix remodeling in outflow tissues ^7^. Whereas prostaglandin F2α receptor agonists are pro-inflammatory and reduce IOP, ocular corticosteroid medications used to suppress the immune system for a variety of ocular disorders can elevate IOP, particularly in glaucoma patients ^9,18^. However, all of these treatments are thought to impact outflow cells and immune cells, making conclusions about their relative contributions impossible.

Consistent with the notion of immune cell regulation of IOP, macrophages are found abundantly in outflow tract tissues such as the trabecular meshwork ^11,19^, as well as around Schlemm’s canal and distal vessels ^11,20^; however, the ontogeny of outflow tract macrophages, namely whether they are prenatally derived (i.e., yolk sac or fetal liver) or are from the adult bone marrow ^21,22^, is unknown. This is important because macrophages are phenotypically distinct based on ontogeny and tissue microenvironment, factors which dictate their function ^23,24^. For example, microglia within different layers of the retina have distinct functions in visual processing ^25^. To address this significant gap in knowledge, we used lineage tracing and ontogeny-specific deletion of macrophages in IOP-relevant tissues of transgenic mice to determine their role in outflow function.

## RESULTS

### Resident tissue macrophages (RTMs) in the outflow tract are enriched in the trabecular meshwork

Aqueous humor is produced by the ciliary body and drains through the conventional outflow tract, initially into the trabecular meshwork (TM) and Schlemm’s canal (SC), then downstream to the distal vessels (DV) and ultimately to the venous circulation (**Figure 1A**). While it has been shown that macrophages populate the outflow tract tissues ^11^, their ontogeny and relative distributions are unknown. To investigate tissue microenvironments within the outflow tract, we immunolabeled cross-sections of the outflow tract tissues in wildtype mice (**Figure 1B**). Smooth muscle actin (SMA) helps identify the trabecular meshwork tissues, whereas CD31 enables detection of the SC and DV. Then, to quantify macrophage distribution, we also imaged flat mounts of outflow tissues (**Figure S1A**), revealing that the greatest density of macrophages was around DVs (∼4X) compared to TM and SC (**Figure 1C**). Hence, we conclude that macrophage distributions are distinct throughout the conventional outflow tract.

**Figure 1.**
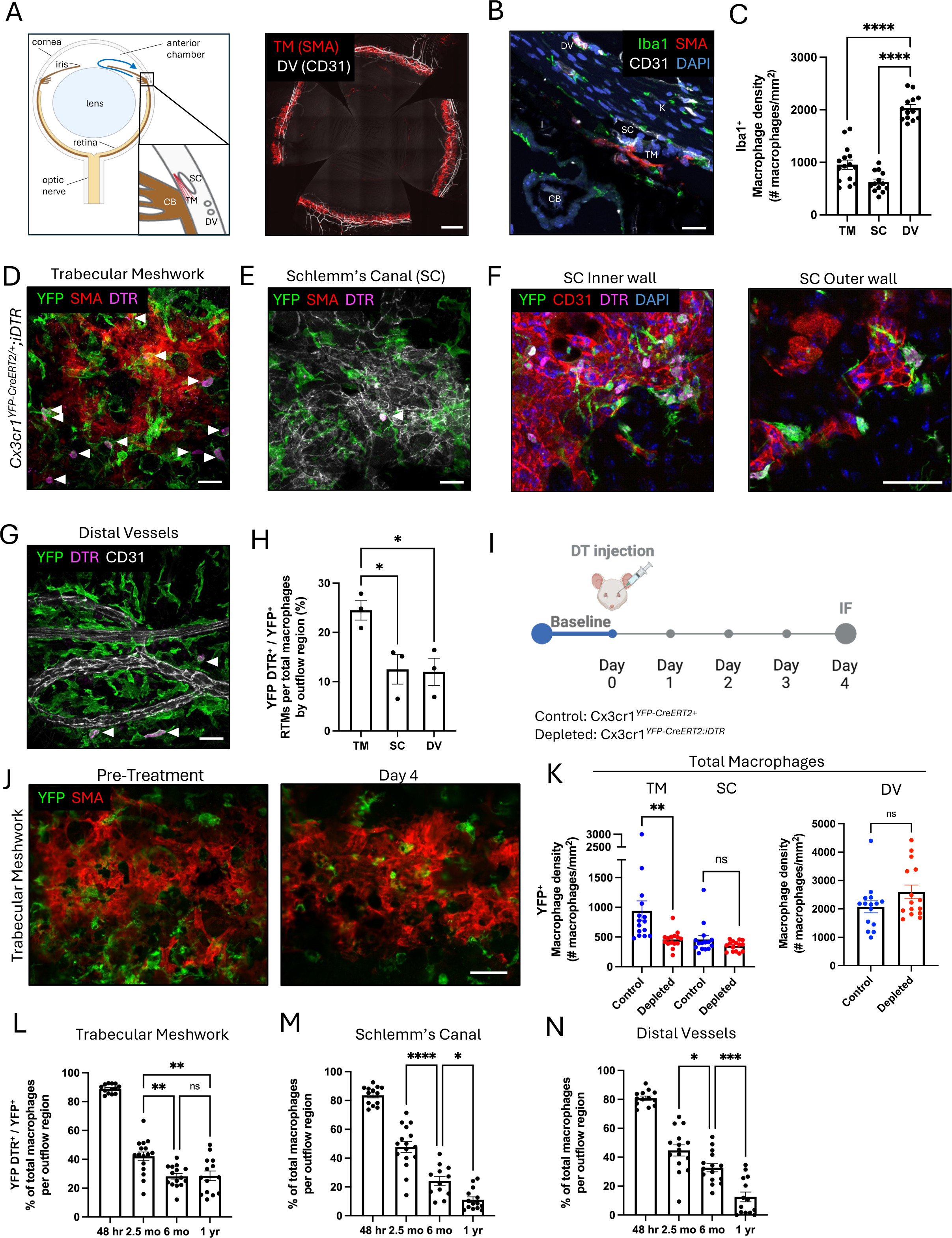
Fate mapping identifies RTMs enriched in the trabecular meshwork within the conventional outflow tract. **(A)** Schematic of the conventional outflow tract in cross section (left). Blue arrow indicates the direction of aqueous flow. En face view of the conventional outflow tract (right) in whole mount anterior segment sections with TM (red) and DVs (white). **(B)** Iba1, SMA and CD31 labeling of outflow tract tissues in an anterior segment cross-section (C57Bl/6 WT). Trabecular meshwork, TM; Schlemm’s canal, SC; distal vessels, DV; cornea, K; ciliary body, CB. Scale bar: 20 µm. **(C)** Density of Iba1^+^ macrophages in TM, SC and DVs in C57Bl/6 WT mice, 5 non-consecutive 40X images per mouse (n=3). **(D)** En face maximum projection images of 11-month-old tam-pulsed *Cx3cr1^YFP-CreERT^*^2^*^/+^;iDTR* mice with RTMs and short-lived macrophages (YFP^+^DTR^+^ and YFP^+^DTR^-^, respectively) populating the TM tissue niche. Arrows mark DTR^+^ RTMs. Scale bar: 25 µm. **(E)** En face images of RTMs and short-lived macrophages in SC of an 11-month-old tamoxifen pulsed *Cx3cr1^YFP-CreERT^*^2^*^/+^;iDTR* mouse. **(F)** RTMs and short-lived macrophages within inner wall (left) and outer wall SC (right) in 3-month-old tam-pulsed *Cx3cr1^YFP-CreERT^*^2^*^/+^;iDTR* mice. Scale bar: 50 µm. **(G)** En face maximum projection images of 11-month-old tamoxifen pulsed *Cx3cr1^YFP-CreERT^*^2^*^/+^;iDTR* mice with RTMs and short-lived macrophages (YFP^+^DTR^+^ and YFP^+^DTR^-^, respectively) surrounding DV. **(H)** Percentage of RTMs in DV, SC and TM from 11-month-old tamoxifen pulsed *Cx3cr1^YFP-CreERT^*^2^*^/+^;iDTR* mice. Each point represents averaged values (5 non-consecutive 40X images) from one mouse (n = 3). **(I)** Schematic of RTM depletion experimental design. IF: immunofluorescence. **(J)** En face maximum projection of Iba1^+^ macrophages at baseline and with depletion in TM. Scale bar: 50 µm. **(K)** Total macrophage density in DV, SC and TM at 4 days following DT treatment (n = 3). **(L-N)** Percent RTMs in TM, SC and around DVs in tam-pulsed *Cx3cr1^YFP-CreERT^*^2^*^/+^;iDTR* tissues collected at 48 hours, 2.5 months, 6 months and 1 year (n = 3 mice per timepoint). Mean ± SEM; one-way ANOVA, Tukey’s post hoc test. *: P < 0.035, **: P < 0.002, ***: P < 0.0002, ****: P < 0.0001, ns: not significant.

We next characterized the ontogeny of identified macrophages in the outflow tract with respect to their distributions. We performed fate mapping of long-lived resident tissue macrophages (RTMs), as these resident macrophages are widely recognized for their homeostatic functions ^26,27^. We used *Cx3cr1^YFP-CreERT^*^2^*^/+^;iDTR* mice ^28,29^ that were tamoxifen pulsed at 6-10 weeks (to induce expression of DTR) and terminated ∼9 months later at 11 months of age. As a control, we examined age-matched *Cx3cr1^YFP-CreERT^*^2^*^/+^;iDTR* mice without tamoxifen treatment (**Figure S1B**). To determine the relative abundance of macrophages by ontogeny across the outflow tract tissues, we immunolabeled for yellow fluorescent protein (YFP) and diphtheria toxin receptor (DTR) ^25^. In this setting, YFP labels 86% of Iba1^+^ macrophages (**Figure S1C, D**), whereas DTR selectively labels long-lived RTMs. We first investigated the relative abundances of total macrophages by immunostaining outflow tract tissues for YFP^+^ cells. Corroborating our analysis with wildtype mice, our results showed the greatest density of macrophages (total YFP^+^ cells) around DVs compared to the TM and SC (**Figure S1E)**. Interestingly, we found that DTR^+^ macrophages (i.e. RTMs) were well represented in the TM, displaying ramified morphologies and characteristic perinuclear distribution of DTR (**Figure 1D**) ^25^. By contrast, DTR^+^ macrophages were less abundant in SC, although ramified morphologies were likewise observed (**Figure 1E**). Moreover, because the inner wall SC (adjacent to TM) and outer wall SC cells are molecularly distinguishable ^30^, we assessed SC macrophages separately using *Cx3cr1^YFP-CreERT^*^2^*^/+^;iDTR* mice at 3 months of age. We found both RTMs and short-lived macrophages in the inner and outer walls of SC (**Figure 1F**). Lastly, we analyzed macrophages around the DVs, finding a low representation of DTR^+^ macrophages that were elongated in appearance (**Figure 1G**). In summary, the TM harbored a greater proportion of RTMs (24.5±2.0%) compared to SC (12.5±3.0%) and DVs (12.0±2.8%; *p* < 0.03) (**Figure 1H**). These results demonstrate that macrophages in the outflow tract have unique spatial distributions, with RTMs being enriched in the TM.

To validate their localization, we depleted RTMs in the outflow tract of *Cx3cr1^YFP-CreERT^*^2^*^/+^;iDTR* mice that were tamoxifen pulsed at 6-10 weeks of age, by subconjunctival injection of diphtheria toxin (DT) at 3 months. As controls, unsensitive tamoxifen treated *Cx3cr1^YFP-CreERT^*^2^*^/+^*mice received DT injections. Four days after DT administration, we harvested tissue and quantified macrophage numbers by outflow tract region (**Figure 1I**). DT treatment resulted in a 58% reduction (*p* < 0.002) in total macrophages in the TM, while numbers were unchanged in SC and DV (**Figure 1J-K**), corroborating our previous results showing RTMs are enriched in TM.

As the persistence of RTMs can vary by tissue ^31^, we next analyzed the longevity and spatiotemporal dynamics of macrophages in the outflow tract. We again applied *Cx3cr1^YFP-CreERT^*^2^*^/+^;iDTR* mice ^28,29^ which were tamoxifen-pulsed at 6-10 weeks and outflow tract tissues collected at 48 hours (to assess baseline CreERT2 recombination), 2.5 months, 6 months and 1 year. In the TM, a steady state population of RTMs (∼25% of total macrophages) was established at 6 months and maintained to 1 year (**Figure 1L**). In contrast, there was a continual decline in the percentage of RTMs in SC and DV over time, until only ∼10% remained at 1 year (**Figure 1M, N**). These results show that macrophage kinetics vary by outflow tract region with only the TM maintaining the highest proportion of RTMs that are stable out to one year.

### Monocyte-derived macrophages and the dual origin of macrophages in the outflow tract

We next characterized the spatial distribution of short-lived macrophages in the outflow tract using the *Ms4a3^Cre^;Rosa^TdTom^*mouse ^32^. The tdTomato (TdTom) fluorescence protein labels monocytes (and granulocytes) but not long-lived prenatal-derived RTMs. Outflow tract tissues from these mice aged 2.5 - 3 months were then evaluated (**Figure 2A**). Short-lived macrophages, identified as Iba1^+^ tdTom^+^, were more abundant around DVs (**Figure 2B**) and comprised ∼50% of total macrophages in the outflow tract (47%, 48% and 55% in TM, SC and DV, respectively) (**Figure S3A**), which corroborates with the spatiotemporal dynamics of macrophages at 2.5 months.

**Figure 2.**
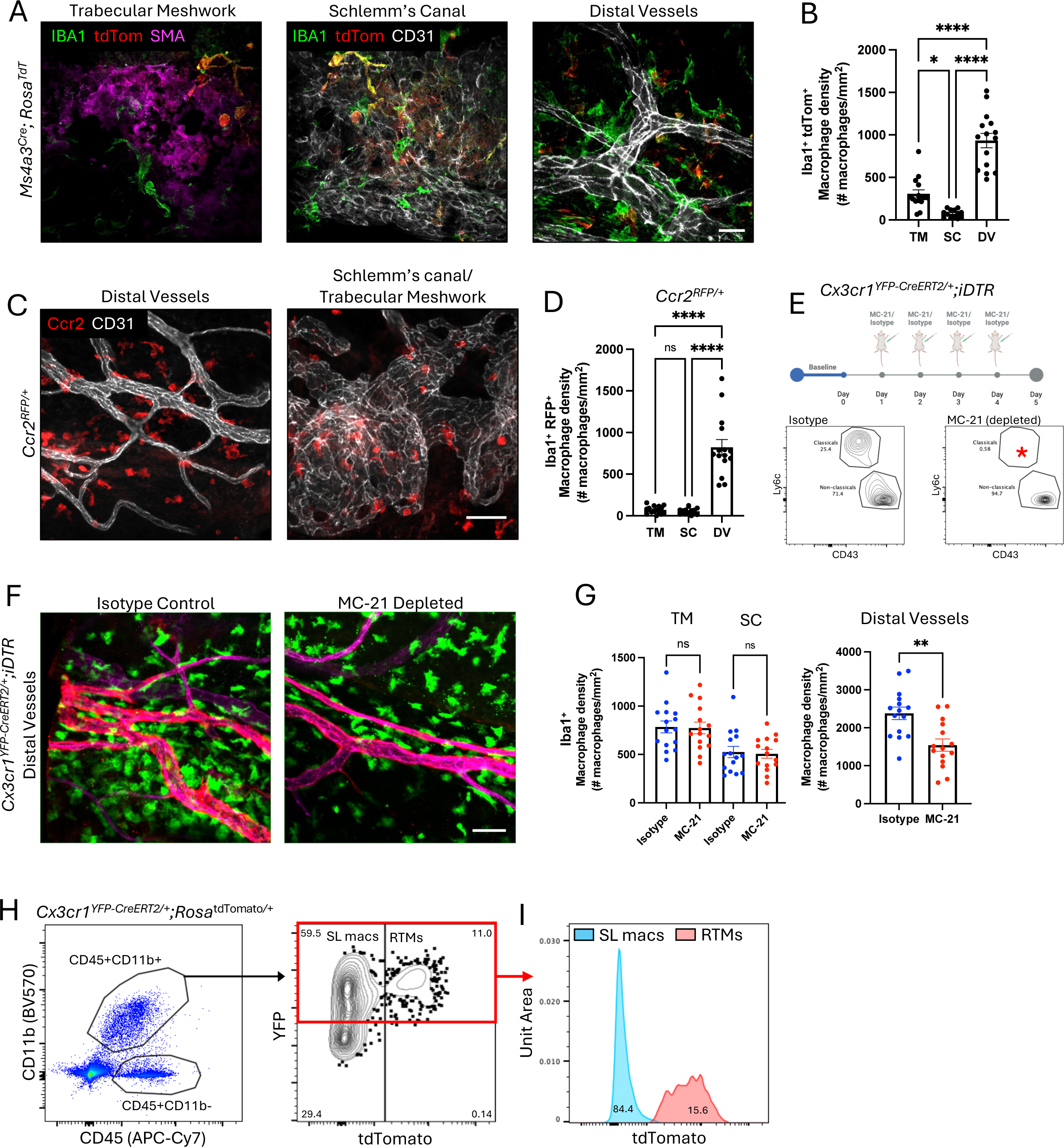
Characterization of short-lived macrophages in the conventional outflow tract. **(A)** En face max projection images of TM, SC and DV from 3-month-old *Ms4a3^Cre^-Rosa^TdTom^*mice show short-lived bone-marrow derived (Iba1^+^ Tdtom^+^) macrophages in outflow tract tissues. Scale bar: 25 µm. **(B)** Quantification of Ms4a3 labeled short-lived macrophage density in outflow tract tissues from 3-month-old *Ms4a3^Cre^-Rosa^TdTom^* mice (n = 3). **(C)** En face max projection images of Ccr2^+^ short-lived macrophages in outflow tract tissues from 3-month-old *Ccr2^RFP/+^* mice. Scale bar: 50 µm. **(D)** Quantification of Ccr2^+^ short-lived macrophage density in DV, SC and TM (n = 3). **(E)** Experimental design with intraperitoneal injections of MC-21 or isotype administered at days 1-4 (top). Validation of monocyte depletion in blood (on day 5) from *Cx3cr1^YFP-CreER/+^;iDTR* mice treated with MC-21 versus isotype using flow cytometry. Classical monocytes were identified as CD43^hi^. Asterisk indicates depleted classical monocyte population (bottom). **(F)** En face maximum projection images of YFP, SMA and CD31 staining of distal vessels with MC-21 and isotype treatment. Scale bar: 50 µm. **(G)** Quantification of total macrophage density in DV, SC and TM in MC21 and isotype treated eyes (n = 3). **(H)** Cells from limbal tissue of 1 year old *Cx3cr1^YFP-CreERT^*^2^*^/+^;Rosa^tdTom/+^* mice. RTMs (GFP^+^ tdTom^+^) and short-lived (SL) (GFP^+^ tdTom^-^) macrophages were identified (n = 8). **(I)** Histograms of RTMs (tdTom^+^; red) and SL macrophages (tdTom^-^; blue) with mean fluorescent intensity. Mean ± SEM; one-way ANOVA, Tukey’s post hoc test (B, D, G). *: P < 0.035, ****: P < 0.0001, ns: not significant.

As a complementary method, we determined the distribution of bone marrow-derived Ccr2^+^ macrophages as C-C motif chemokine receptor 2 (CCR2) is needed for the egress of monocytes from the bone marrow ^28^. In *Ccr2^RFP/+^*mice, Ccr2^+^ macrophages are Iba1^+^ RFP^+^. Ccr2^+^ macrophages were more abundant around DVs compared to TM and SC (**Figure 2C, D**). In the combined SC/TM maximum projection images (**Figure 2D**), the TM (not labeled) is located below the plane of SC (CD31). We note that Ccr2 expression is downregulated when monocytes fully differentiate into macrophages in the tissue ^33^, thus underestimating the number of total short-lived macrophages.

Using another method, we pharmacologically targeted Ccr2 with the Ccr2 antibody, MC-21 ^34^, to deplete blood classical monocytes. *Cx3cr1^YFP-CreERT^*^2^*^/+^;iDTR* mice were treated with MC-21 or isotype control by intraperitoneal injection daily for 4 days (**Figure 2E**). Peripheral blood mononuclear cells were isolated, and flow cytometry demonstrated efficient depletion of classical monocytes in MC-21 treated mice (**Figure 2E**). Consistent with our previous results that RTMs are enriched in the TM, our analysis of the outflow tract tissues following MC-21 treatment revealed a reduction in macrophage numbers around DVs but were unchanged in TM and SC (**Figure 2F, G)**. These results are congruent to the distribution of short-lived macrophages in *Ccr2^RFP/+^* and *Ms4a3^Cre^-Rosa^TdTom^*mice. In short, outflow tract macrophages are of dual origin with the greatest number of monocyte-derived macrophages around DVs.

Lastly, we sought to characterize both macrophage ontogenies by flow cytometry. We used *Cx3cr1^YFP-CreERT^*^2^*^/+^;Rosa*^tdTom*/+*^ mice that were tamoxifen pulsed at 6-10 weeks and subsequently aged 1 year post tamoxifen ^35^ (**Figure S2A)**. Multi-dimensional flow cytometry of dissociated cells from outflow tract tissues was used with our previously established panel ^25^, which included CD45, CD11b, Ly6C, F4/80, Ly6G, CD64, I/A-I/E, CD11c, and endogenous YFP and tdTom reporters. Our results showed that 15.6% of macrophages were YFP^+^ tdTom^+^ RTMs, whereas 84.4% were YFP^+^ tdTom^-^ short-lived (**Figure 2H**) which corroborates with the spatiotemporal dynamics of RTMs at 1 year. t-Stochastic Neighbor (tSNE) analysis shows that marker expression was distinct between RTMs and short-lived macrophages (**Figure S2B, C**) on the basis of higher CD64 expression (*p* < 0.0001) by RTMs versus higher Ly6c expression (*p* < 0.002) by short-lived macrophages (**Figure S2D**).

### RTM-specific role in outflow homeostasis, associated with extracellular matrix turnover in the trabecular meshwork

RTMs have tissue-specific functions in homeostasis, as widely appreciated with microglia, osteoclasts and alveolar macrophages ^36^. To test whether RTMs have a functional role in outflow homeostasis, we depleted RTMs in the outflow tract of tamoxifen pulsed *Cx3cr1^YFP-CreERT^*^2^*^/+^;iDTR* mice and *Cx3cr1^YFP-CreERT^*^2^*^/+^*control mice at 3 months of age using DT as before. At 4 days post-DT treatment, significant depletion of DTR^+^ RTMs was detected in the TM/SC (**Figure 3A**) and DV (depleted by 98% and 77%, *p* < 0.002 and *p* < 0.0001, respectively; **Figure 3B**). To interrogate the function of RTMs in conventional outflow homeostasis, we measured IOP in RTM depleted and control mice at baseline and 4 days post-treatment. RTM-depleted eyes showed a significant increase in IOP (18.4±0.4 mmHg) compared to their baseline readings (16.9±0.3 mmHg, *p* < 0.0002), as well as compared to IOPs of *Cx3cr1^YFP-CreERT^*^2^*^/+^* control mice (16.1±0.3 mmHg, *p* < 0.002) (**Figure 3C**). In the conventional outflow tract, most of the outflow resistance is generated by the proximal portion (TM and SC) of the outflow pathway (**Figure 3D**). To assess whether RTMs affect IOP at the level of the TM and SC, we measured outflow facility ^37^, which quantifies pressure-dependent movement of aqueous humor out of the eye ^38^. Facility specifically through the proximal outflow tract was assessed, as vascular tone of the distal outflow vessels is lost with *ex vivo* measurements. Corresponding to an elevated IOP, results show that outflow facility was significantly reduced by 59% in eyes where RTMs were depleted compared to control eyes (1.0±0.1 versus 1.7±0.2 nl/min/mmHg, *p* < 0.002) (**Figure 3E**). Hence, these data suggest that TM-enriched RTMs function in IOP homeostasis at the level of the proximal outflow pathway, the site of pathology responsible for ocular hypertension in glaucoma.

**Figure 3.**
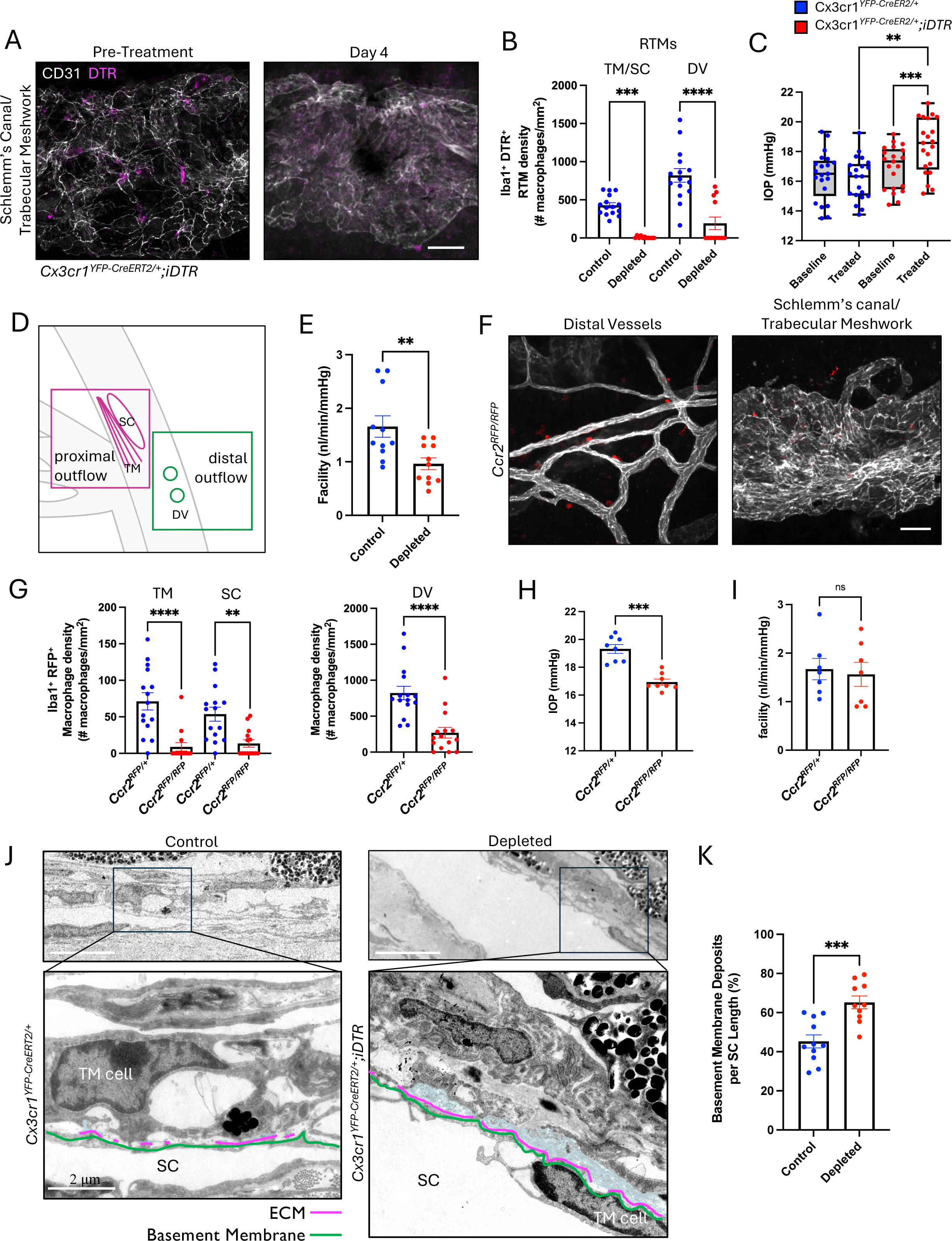
RTMs contribute to IOP homeostasis by affecting conventional outflow, associated with extracellular matrix turnover. **(A)** En face maximum projection images of RTM depletion 4 days after subconjunctival DT injection in tam-pulsed *Cx3cr1^YFP-CreERT^*^2^*^/+^;iDTR* mice. Scale bar: 50 µm. **(B)** RTMs in TM/SC and DV following DT treatment (n = 3). **(C)** Intraocular pressure with RTM depletion compared to control eyes at baseline and after DT. Each point represents the average of paired eyes (n = 21/group from 2 experiments). **(D)** Schematic differentiating the proximal and distal regions of the conventional outflow tract. **(E)** Outflow facility measured *ex vivo* in RTM depleted and control eyes (n = 11 per group). Each point represents the average of paired eyes. **(F)** Max projection images of depleted Ccr2*^RFP/RFP^* around DVs (left) and in SC/TM (right) of 3-month-old *Ccr2^RFP/+^* mice. Scale bar: 50 µm. **(G)** Ccr2^+^ macrophages in TM, SC and DV in *Ccr2^RFP/+^* and *Ccr2^RFP/RFP^* mice (n = 3). **(H)** IOP in control and short-lived macrophage depleted eyes. Each point represents the average of paired eyes (n = 8). **(I)** Outflow facility in eyes of control *Ccr2^RFP/+^*and *Ccr2^RFP/RFP^* mice (n = 8, each point is the average of paired eyes). **(J)** Transmission electron micrographs of the inner wall of SC, underlying basement membrane (green) and ECM material (blue) in control and RTM depleted eyes at 2500X and 12,000X mag (Pink lines show basement membrane overlaid with ECM material). **(K)** Percentage of inner wall SC basement membrane covered by ECM material in RTM-depleted (*Cx3cr1^YFP-CreERT^*^2^*^/+^;iDTR*) and control eyes (*Cx3cr1^YFP-CreERT^*^2^*^/+^*) (n = 8 per group). Each point represents percent of basement membrane deposits over the SC length in a single cross-section. (n=8 mice/group, 2-3 cross-sections taken from different quadrants). Mean ± SEM; one-way ANOVA, Tukey’s post hoc test (B, C, G) and unpaired Student’s two-tailed t test (E, H, I, K). *: P < 0.035, **: P < 0.002, ***: P < 0.0002, ****: P < 0.0001, ns: not significant.

To assess whether this role for RTMs in conventional outflow homeostasis is ontogeny-specific, we next depleted short-lived macrophages and assessed measures of outflow function. We took advantage of hemizygous *Ccr2^RFP/+^* versus homozygous *Ccr2^RFP/RFP^* knockin mice, which we confirmed has a reduction in monocyte recruitment to the outflow tract tissues (**Figure 3F, G**). Interestingly, *Ccr2^RFP/RFP^*mice did not show an increase in IOP (16.9±0.2 mmHg) compared to *Ccr2^RFP/+^* controls (19.3±0.3 mmHg) (**Figure 3H**). As well, our results show that *Ccr2^RFP/RFP^* eyes exhibited no effect on outflow facility (**Figure 3I**), i.e. resistance of the proximal outflow tract. As a separate approach, we impaired monocyte-derived macrophages via MC-21 treatment. MC-21 also did not show an increase in IOP (15.9±0.9 mmHg) compared to baseline (17.0±0.8 mmHg), and there was no effect on outflow facility (**Figure S3C, D)**, like in *Ccr2^RFP/RFP^* mice. These results suggest that monocyte-derived macrophage depletion does not lead to an increase in IOP, which is in contrast with what we observed with RTM depletion. Our data with short-lived macrophage depletion rule out an effect on the primary regulator of IOP, the TM and SC, which is specifically governed by RTMs.

One of the major mechanisms that regulates outflow resistance in the TM and SC and is dysregulated in glaucoma is the turnover rate of extracellular matrix (ECM) ^39,40^. Like TM cells, macrophages are widely appreciated participants in ECM turnover ^41^. Thus, we asked if RTM depletion affects ECM content in the conventional outflow tract. To address this question, we quantified the amount of basement membrane deposits underlying the inner wall of SC, which is an established surrogate measurement of ECM turnover in the resistance-generating region between the TM and SC ^42,43^. Basement membrane deposits were quantified across the full length of inner wall of SC in anterior segment cross-sections, with 2-3 cross-sections taken from different quadrants per mouse eye (n = 8 mice per group) in RTM-depleted (DT treated *Cx3cr1^YFP-CreERT^*^2^*^/+^;iDTR*) and control (DT treated *Cx3cr1^YFP-CreERT^*^2^*^/+^*) groups. Using high magnification transmission electron micrographs of the TM and SC, RTM-depleted eyes contained significantly greater ECM deposits covering the inner wall of SC compared to controls (65.2±3.3% versus 45.3±3.3% of inner wall of SC length, respectively; *p* < 0.0002) (**Figure 3J, K; S4**). By contrast, when we compared monocyte-derived macrophage depleted *Ccr2^RFP/RFP^* and control *Ccr2^RFP/+^* mice, we found no difference in the amount of basement deposits in short-lived macrophage depleted versus control mice (25.4±2.1 versus 29.5±2.9%, respectively; p = 0.28), consistent with functional measurements (**Figure S5A, B**). In summary, RTMs have a specific role in IOP homeostasis associated with ECM turnover, a function that is not extended with short-lived macrophages.

## DISCUSSION

The role of the immune system in IOP homeostasis is unknown. Using complementary mouse model systems to dissect out relative roles, we show that RTMs in the conventional outflow pathway are essential to maintain a healthy IOP, and that RTMs and short-lived macrophages have differential effects on IOP regulation. Mechanistically, we found that TM-enriched RTMs affect outflow resistance by altering ECM turnover dynamics. In contrast, short-lived macrophages are more abundant around DVs and have no apparent role in the regulation of outflow resistance and ECM abundance in the TM in health.

Our studies reveal that macrophages in the outflow tract are phenotypically heterogenous. Among different organs and tissues within the body, macrophages demonstrate heterogeneity in ontogeny, e.g., prenatally derived RTMs reside in organs such as the brain and skin, whereas bone-marrow derived monocytes maintain macrophages in the gut and heart ^31^. In the eye, macrophages that are located in the retina are largely comprised of microglia ^44^, whereas macrophages in the cornea and choroid originate from bone-marrow derived monocytes ^25^. In contrast, the conventional outflow tract is populated by both RTMs and monocyte-derived macrophages, like iris/ciliary body and optic nerve tissues ^25^. While there is a small population of RTMs, the SC and distal outflow tract (DV) can be considered an “open” niche ^31^ with the ability to recruit monocyte-derived macrophages to the outflow tract tissues via access to the systemic venous blood circulation. In contrast, the TM is positioned behind the blood-aqueous barrier formed by tight junctions of the inner wall of SC ^45^, where RTMs are enriched. In addition to heterogeneity by ontogeny, outflow tract macrophages show differential expression of myeloid cell markers by multi-dimensional flow cytometry.

Significantly, we demonstrate for the first time a direct role for the immune system in normal IOP homeostasis at steady state, which contrasts with prior work showing indirect, correlative evidence. For example, previous studies show that laser trabeculoplasty of the TM, which alters the steady-state condition, results in temporary IOP reduction due to decreased outflow resistance, with an associated influx of mononuclear phagocytes. This led the authors to hypothesize that mononuclear cells modulate IOP ^8^. Interestingly, our study showed that depletion of short-lived macrophages did not increase IOP in a steady state setting, and had no effect on outflow resistance. This discrepancy between the role of short-lived macrophages in the TM during laser challenge and at steady state needs to be resolved in a controlled experimental setting.

RTMs have known homeostatic functions in many tissue niches within the body ^36^. Macrophages can also perform homeostatic functions in tissues that are predominately populated by monocyte-derived macrophages such as the intestine ^46^ and aorta ^47^. Uniquely, the conventional outflow tract is populated by both RTMs and short-lived macrophages that may have differing roles in IOP regulation in the healthy state. This affords the potential to therapeutically target specific macrophage subpopulations in the outflow tract to lower IOP.

Macrophages are also likely important in glaucoma pathogenesis, the second leading cause of blindness in the world. As such, pathway analyses of genome-wide association data identified macrophage relevant biological processes such as “phagocytosis engulfment” in glaucoma patients ^12^. Moreover, histological examination of postmortem human eyes with glaucoma showed increased CD163^+^ macrophages in glaucomatous optic nerves and variable distribution of CD163^+^ cells in the outflow tract compared to eyes without glaucoma ^48^. Future studies are needed to investigate the roles of RTMs and short-lived macrophages in the human outflow tract in normal IOP homeostasis versus disease.

## ACKNOWLEDGEMENTS

This study was funded by K08EY032202 (K.C.L.), Research to Prevent Blindness CDA (K.C.L.), Duke Strong Start Award (K.C.L.), R01EY030906 (D.R.S.), U01EY034687 (D.R.S.), R01EY022359 (W.D.S.), R01EY028608

(W.D.S.), R01EY030124 (W.D.S.), BrightFocus Foundation (W.D.S.), P30EY005722 (Duke Eye Center), and Research to Prevent Blindness (Duke Eye Center). We thank Y. Hao for TEM (Duke Eye Center), E. Miao (Duke University) for gifting of *Ccr2^RFP/RFP^* mice, and O. Kolupev (Duke Eye Center) for assistance with flow cytometry analysis.

## AUTHOR CONTRIBUTIONS

K.C.L. designed, performed and analyzed experiments and wrote the manuscript. A.O.G and D.S. performed and analyzed experiments and performed animal husbandry. M.F.S., M.L.D.I., R.A.K., R.M., J.K., V.B. and R.B. performed and analyzed experiments. M.M. and F.G. contributed to data analysis and interpretation and manuscript writing. S.W.M.J. contributed to project design, data analysis and interpretation and manuscript writing. W.D.S. and D.R.S. were involved in all aspects of the project and wrote the manuscript.

## SUPPLEMENTAL INFORMATION

Document S1. Figures S1-S5

## METHODS

### Mice

C57BL/6J mice were purchased from The Jackson Laboratory (Bar Harbor, Maine) as well as *Ms4a3^Cre^-Rosa^TdTom^* mice [stock No. 036382], *Cx3cr1^YFP-CreERT^*^2^ *^YFP-CreERT^*^2^ mice [stock No. 021160], and the inducible DTR mouse line, *Rosa26^R26R-DTR/R26R-DTR^* [stock No. 007900]. Rosa-lsl-tdTomato (*R26^RFP^*) line was kindly provided by Dr. M. Dee Gunn (Duke University) and was originally purchased from Jackson Laboratory [stock No. 007914]. *Ccr2^RFP/RFP^* mice were kindly provided by Dr. Edward Miao (Duke University) and were originally purchased from Jackson Laboratory. All mice are housed at a 12-hour light/dark cycle, barrier-free and specific-pathogen-free facility at Duke University Eye Center (Durham, NC). Mice were fed 5001 laboratory rodent diet *ad libitum* (LabDiet). All procedures were approved by the Institutional Animal Care and Use Committee at Duke University, and the procedures were carried out in accordance with the approved guidelines.

### Immunolabeling

Mouse eyes were fixed in 4% paraformaldehyde (PFA) for 30 minutes at 4°C and then 30 minutes at room temperature. Eyes were washed in PBS. The anterior cup was dissected off and cut into 4 quadrants each. Anterior segments were treated with a blocking and permeabilization solution of 5% donkey or goat serum, 1% Triton-X and 0.1% Tween-20. Samples were then treated with solutions containing various primary antibodies such as: CD31 (Millipore, mAb13982), αSMA conjugated to Cy3 (Millipore, C6198), IBA1 (Fujifilm, #019-19741), GFP (in-house), RFP (abcam, ab62341) and DTR (R&D Systems, AF-259-NA) in 5% donkey or goat serum overnight at 4°C. After washing samples in 0.2% Tween-20 in PBS, they were treated with filtered solutions of secondary antibodies and incubated overnight at 4°C. Samples were washed in 0.2% Tween-20 solution and mounted to slides using Immu-Mount solution (Fisher Scientific). Images were taken using a Nikon Eclipse 90i confocal laser-scanning microscope (Melville). 40X Z-stack images of 1.2 µm thickness through trabecular meshwork/Schlemm’s canal or distal vessels were collected.

For cryosections, eyes were enucleated and fixed with 4% PFA overnight at 4°C. PFA was removed, and eyes were washed twice with 1X PBS. Anterior cups were dissected and placed in 30% sucrose overnight at 4°C. Anterior cups were carefully placed in OCT in a 10x10x5 mm cryomold and frozen at -80°C. 12 µm sections were cut using a cryostat and placed on positively charged slides. These slides were stored at -20°C. A hydrophobic barrier pen was used to create a water barrier around the sections. Slides were washed with PBS, adding dropwise to remove OCT. Slides were washed for 2 minutes with 70% ethanol and 2 minutes with 100% ethanol. The tissue was blocked with 10% goat serum in 1% Triton-X solution for 1 hour at room temperature. Tissue was then incubated with primary antibody in blocking buffer overnight at 4°C. Tissue was washed with PBS and incubated with secondary antibodies for 1 hour at room temperature. Samples were washed with PBS and mounted using ProLong Diamond Antifade Mountant with DAPI (Thermo Fisher) and had a coverslip added. Slides were left to dry overnight at room temperature before sealing with clear nail polish and imaging. A negative control sample was always prepared without primary antibody. Images were taken using a Nikon Eclipse 90i confocal laser-scanning microscope (Melville). Z-stack images of 1 µm thickness were taken of each section and converted into maximum projection images.

### Tissue Harvesting

For flow cytometry, outflow tract tissues (TM, SC, DV) were harvested from PBS cardiac perfused and freshly euthanized mice. Limbal strips were dissected from anterior segments after the removal of the iris and ciliary body tissue. Samples were digested in 1X collagenase D with shaking at 450rpm at 37°C for 30 minutes. Pellet was resuspended in papain digestion solution (Worthington Biochemical) with DNase for an additional 30 minutes with shaking. Digestion was then halted with an ovomucoid inhibitor solution. Pellet was triturated and washed and resuspended in HBSS. Solution was passed through a 40µm cell strainer and resuspended in FACS+EDTA buffer. Antibodies were added to cell solutions in FACS+EDTA buffer and incubated for 25 minutes on ice. Cells were resuspended in 500uL viability dye and incubated covered on ice. Final wash was conducted using 3mL FACS+EDTA buffer and remaining cells were treated with FACS stabilizing fixative. Cells were stored at 4°C for up to 24 hours before analysis.

### Flow Cytometry Immunolabeling

The flow cytometry panel and immunolabeling panel was previously established ^44^. Single cell suspensions of limbal tissue were incubated in 5% normal mouse serum, 5% normal rat serum, 1% Fc Block (eBiosciences, San Diego, CA) and a combination of fluorophore-conjugated primary antibodies against CD45 (BioLegend, 103116), Ly6C (BioLegend, 128018), Ly6G (BioLegend, 127622), CD64 (BioLegend, 139309), CD11c (BioLegend, 117336), CD11b (Biolegend, 101233), F4/80 (BioLegend, 123115), and I-A/I-E (BD Biosciences, 563415). Samples were stained on ice for 25 minutes. Cells were treated with Aqua Live/Dead viability dye (Thermo Fisher Scientific, Waltham, MA) in PBS for 15 minutes on ice. Samples were then washed and fixed with 0.4% paraformaldehyde in PBS. BD Fortessa flow cytometer and BD FACSDiva software (BD Biosciences) were used to acquire data. Raw flow cytometry data was analyzed using FlowJo software (FlowJo LLC, Ashland, OR).

### Injections

Tamoxifen (20 mg/mL; Sigma-Aldrich) was dissolved in corn oil. 5-6-week-old mice received 2 intraperitoneal injections of 75 mg/kg tamoxifen with one day between injections.

Diphtheria toxin (0.5 ng/µl; Sigma-Aldrich) at a volume of 10 µl was injected into the subconjunctival space of anesthetized mice 4 days prior to IOP, facility measurements and euthanasia.

Mice received intraperitoneal injections of 20 µg of anti-CCR2 antibody (MC-21) ^34^ or isotype antibody (Bio X Cell, B0086) once daily for 4 days prior to IOP, euthanasia and outflow facility measurements.

### Intraocular Pressure Measurement

Mice were anesthetized with isoflurane (2% v/v for induction; 1.0%–1.5% v/v for maintenance using a mouse nose cone) and IOP was measured by rebound tonometry using the iCare Tonolab tonometer (iCare, Finland) approximately 5 minutes following induction of anesthesia. IOPs are an average of 6 measurements. The time of day of IOP measurements were consistent within experiments (9:00 am - 12:00 pm).

### Outflow Facility Measurement

Mice were euthanized by isoflurane inhalation, followed by decapitation as the secondary method. Eyes were enucleated and immediately perfused. Each eye was mounted on a stabilization platform in a perfusion chamber using a small amount of cyanoacrylate glue (Loctite, Westlake, OH). The perfusion chamber was filled with D-glucose in PBS (DBG) at 35°C. A beveled, sharpened glass microneedle filled with DBG was inserted into anterior chamber using a micromanipulator. Outflow facility was measured with the iPerfusion system ^37^. After cannulation, the eyes underwent a 30-minute acclimation at 12 mmHg, before starting the pressure steps to measure facility. The steps started at 5 mmHg and increased sequentially 1.5 mmHg each step until reaching 17 mmHg, then dropping to 8 mmHg for the final step.

Data analysis was carried out as described previously ^37^. Briefly, a nonlinear flow-pressure model was used to account for the pressure dependence of outflow facility in mice. The treatment groups were randomized and masked to the perfusionist.

### Transmission Electron Microscopy

After enucleation, whole eyes were fixed in 2% PFA + 2% glutaraldehyde in PBS for 1 hour at room temperature and then overnight at 4°C. Eyes were washed twice with PBS, immersed in 2% osmium tetroxide in 0.1% cacodylate buffer, dehydrated, and embedded in Epon 812 resin. For light microscopy, semi-thin sections of 0.5 µm were prepared and counterstained with 1% methylene blue. For electron microscopy, 65-75 nm thin sections were collected on a Leica EM CU7 and counterstained with a solution of 2% uranyl acetate and 3.5% lead citrate. Samples were examined using a JEM-1400 transmission electron microscope at 60 kV. An Orius 1000 charge-coupled device camera was used to collect images.

### Macrophage Quantification

Using 40X z-stacks of TM/SC or DVs, macrophages were quantified in ImageJ. A 4x4 grid was overlaid on the image to facilitate quantification. Macrophages on the same plane as SC (identified with CD31) but not in TM (identified with αSMA) were counted as macrophages on SC. Macrophages on the same plane as trabecular meshwork (αSMA) but not in SC or ciliary body were counted as macrophages in TM. In distal vessel z-stacks, macrophages in direct contact with DVs were counted as DV macrophages. The SC or DV areas (CD31) were measured using maximum projection images. TM, SC and DV macrophage densities were calculated as macrophage number divided by the area of SC (for TM and SC) or distal vessels. Macrophage densities were quantified in every other 40X field of view with 5 total images per mouse and 3 mice per average density.

### Basement Membrane Material Quantification

This method has been previously described ^42^. Briefly, TEM images were taken at 12,000X magnification and quantified in ImageJ. Continuous basement membrane material was identified as fine fibrillar and granular material under the inner wall of SC. Basement membrane material was measured along the entire anterior-to-posterior length of SC in each section. The cumulative length of each continuous portion of basement membrane was divided by the total inner wall length for that section, and a percentage was calculated. 2-3 sections taken 90 degrees apart from 8 mice per group were quantified.

### Statistics

Statistical analyses were performed using GraphPad Prism v.8 (GraphPad Software, La Jolla, CA). A P value of 0.05 or less was considered significant and data are presented as mean ± SEM.

---

**Supplemental Figure 1. Macrophages are present throughout the outflow tract tissue of WT C57Bl/6 mice. (A)** En face maximum projection images of Iba1, SMA and CD31 staining in outflow tract tissues (DV, SC, TM) in adult wildtype C57Bl/6 mice. Scale bar: 50 µm. **(B)** En face maximum projection images of YFP, Iba1 and CD31 staining in 3-month-old *Cx3cr1^YFP-CreERT^*^2^*^/+^*mice. Scale bar: 50 µm. **(C)** En face maximum projection images of YFP, Iba1 and CD31 staining in 3-month-old *Cx3cr1^YFP-CreERT^*^2^*^/+^;iDTR* mice. Scale bar: 50 µm. **(D)** Quantification of co-localization of Iba1 and YFP staining in DV and TM/SC (n = 1). 1. **(E)** Density of macrophages that populate the TM, SC and DV normalized to CD31 area. Data were collected from 3 11-month-old tamoxifen pulsed *Cx3cr1^YFP-CreERT^*^2^*^/+^;iDTR* mice. Graphs show mean ± SEM; one-way ANOVA with Tukey’s post hoc test (E). ****: P < 0.0001, ns: not significant.

**Supplemental Figure 2. Multidimensional flow cytometry shows separate macrophage subpopulations from limbal outflow tract tissues. (A)** Cells from limbal tissue of 1-year-old *Cx3cr1^YFP-CreER/+^;Rosa^tdTom/+^*mice were sorted by live/dead signal, gated on singlets, then CD45^+^ and CD11b^+^ macrophages, then RTMs (YFP^+^ tdTom^+^) and short-lived (YFP^+^ tdTom^-^) macrophages were identified (n = 16 pooled eyes from 8 mice). **(B)** Histograms of the markers CD45, Ly6c, CD64, CD11b and YFP in RTMs (tdTom^+^; red) and short-lived (tdTom^-^; blue) macrophages. Numbers on the right indicate mean fluorescent intensity. **(C)** t-stochastic neighbor embedding strategy with clustering of short-lived macrophages (blue) and RTMs (orange) using markers CD45, CD11b, CD64, Ly6c and YFP. **(D)** Bar graphs of mean fluorescence intensity of markers F4/80, Ly6g, D45, CD11b, MHCII, CD11c, Ly6c and YFP in RTMs in short-lived macrophages (YFP^+^ tdTom^-^) and RTMs (YFP^+^ tdTom^+^).

**Supplemental Figure 3. MC-21 depletes monocytes around distal vessels and alters IOP. (A)** Percentage of short-lived macrophages in outflow tract tissues from 3-month-old *Ms4a3^Cre^-Rosa^TdTom^*mice (average of 5 non-consecutive 40X images per mouse, n = 3). **(B)** Validation of monocyte depletion in blood (on day 5) from *Cx3cr1^YFP-CreER/+^;iDTR* mice treated with MC-21 versus isotype using flow cytometry. Single cells were gated on CD45 for leukocytes, CD11b^hi^ and Ly6g^lo^ for myeloid cells, then CD115^hi^ for monocytes, and CD43^hi^ for classical monocytes. Asterisk indicates depleted classical monocyte population. **(C)** IOP of MC-21 and isotype treated eyes at baseline and post-injection (n = 6). **(D)** Outflow facility of MC-21 and isotype treated eye (n = 6). Graphs show mean ± SEM; one-way ANOVA with Tukey’s post hoc test (A), two-tailed paired t-test (C), and d unpaired Student’s two-tailed t test (D). **: P < 0.002, ns: not significant.

**Supplemental Figure 4.** ECM **underling SC basement membrane length in RTM-depleted mice.** Unlabeled transmission electron micrographs of the inner wall of SC, underlying basement membrane and grainy-appearing ECM material in control and RTM depleted-eyes at 2500X and 12,000X magnification.

**Supplemental Figure 5. Short-lived macrophage depletion does not affect extracellular matrix turnover. (A)** Transmission electron micrographs of the inner wall of SC, underlying basement membrane (green) and ECM material (pink) in control and short-lived macrophage depleted eyes at 12,000X magnification. Pink lines show basement membrane overlaid with ECM material. **(B)** Quantification of the percentage of inner wall SC basement membrane covered by extracellular matrix material in short-lived macrophage depleted and control eyes (n = 3 per group, 2 cross-sections per mouse). Graph shows mean ± SEM; unpaired Student’s two-tailed t test (B). ns: not significant.

