## Supplemental Figures for "Resident Tissue Macrophages Govern Intraocular Pressure Homeostasis"

### Supplemental Figure 1

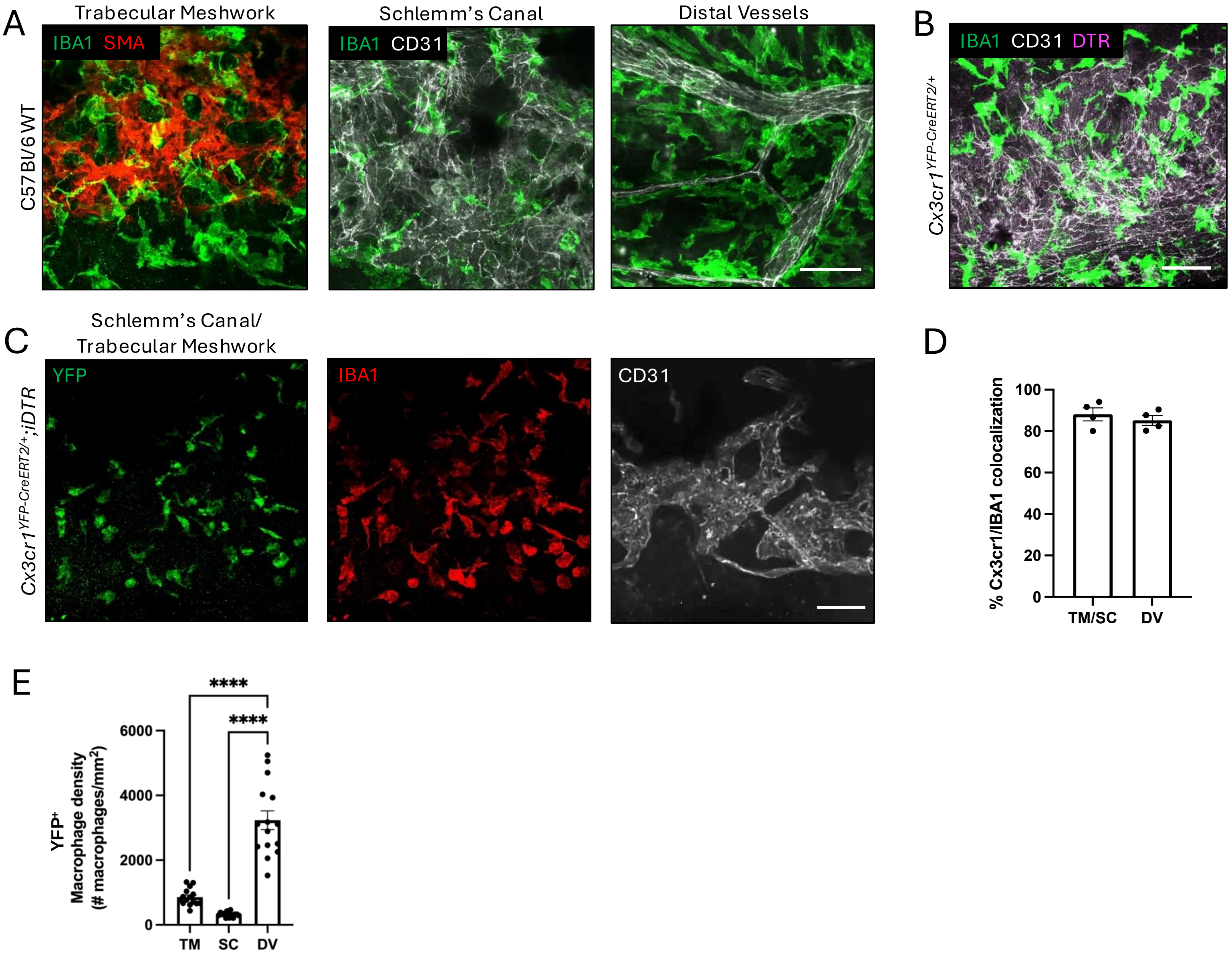

### Supplemental Figure 2

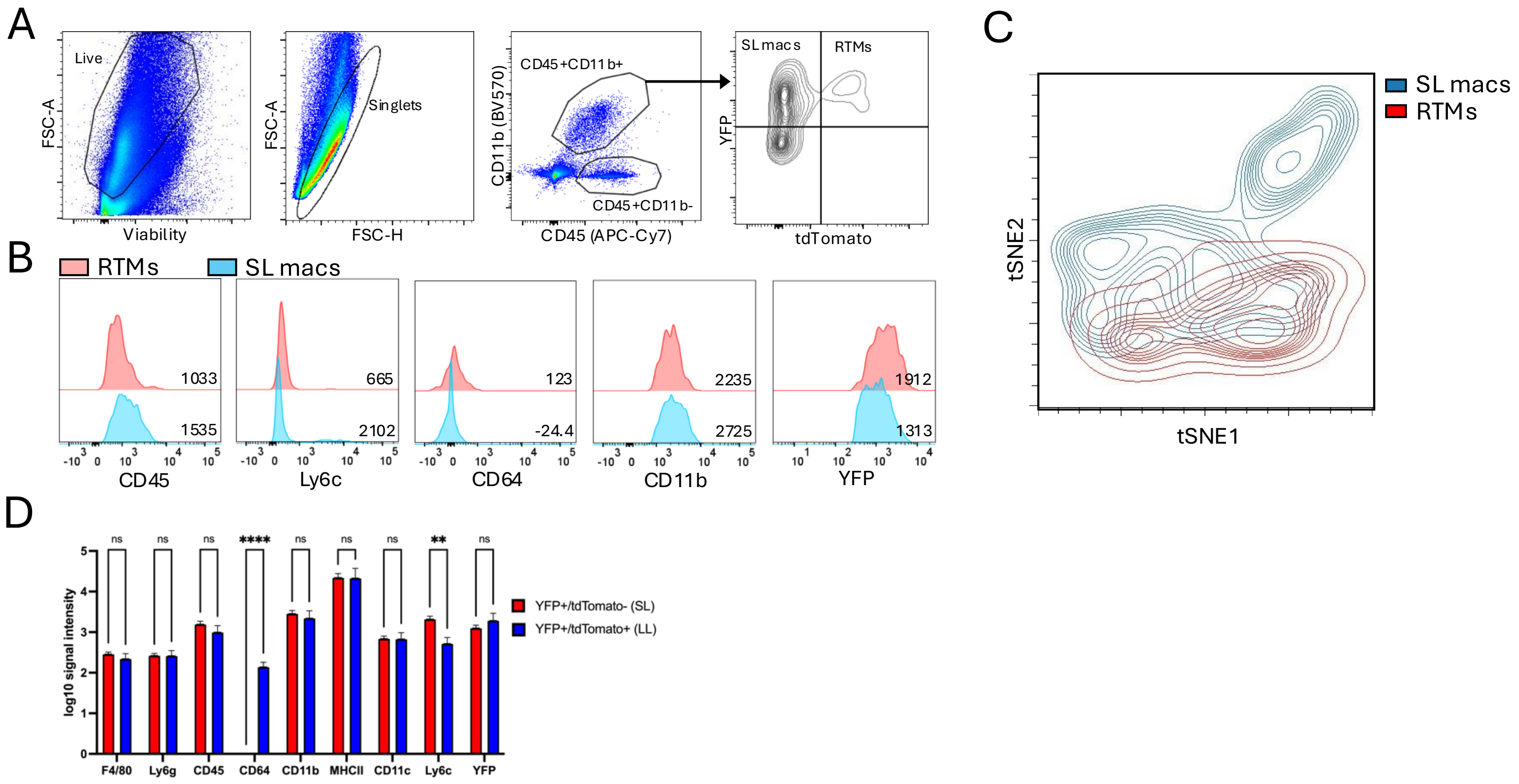

Supplemental Figure 3

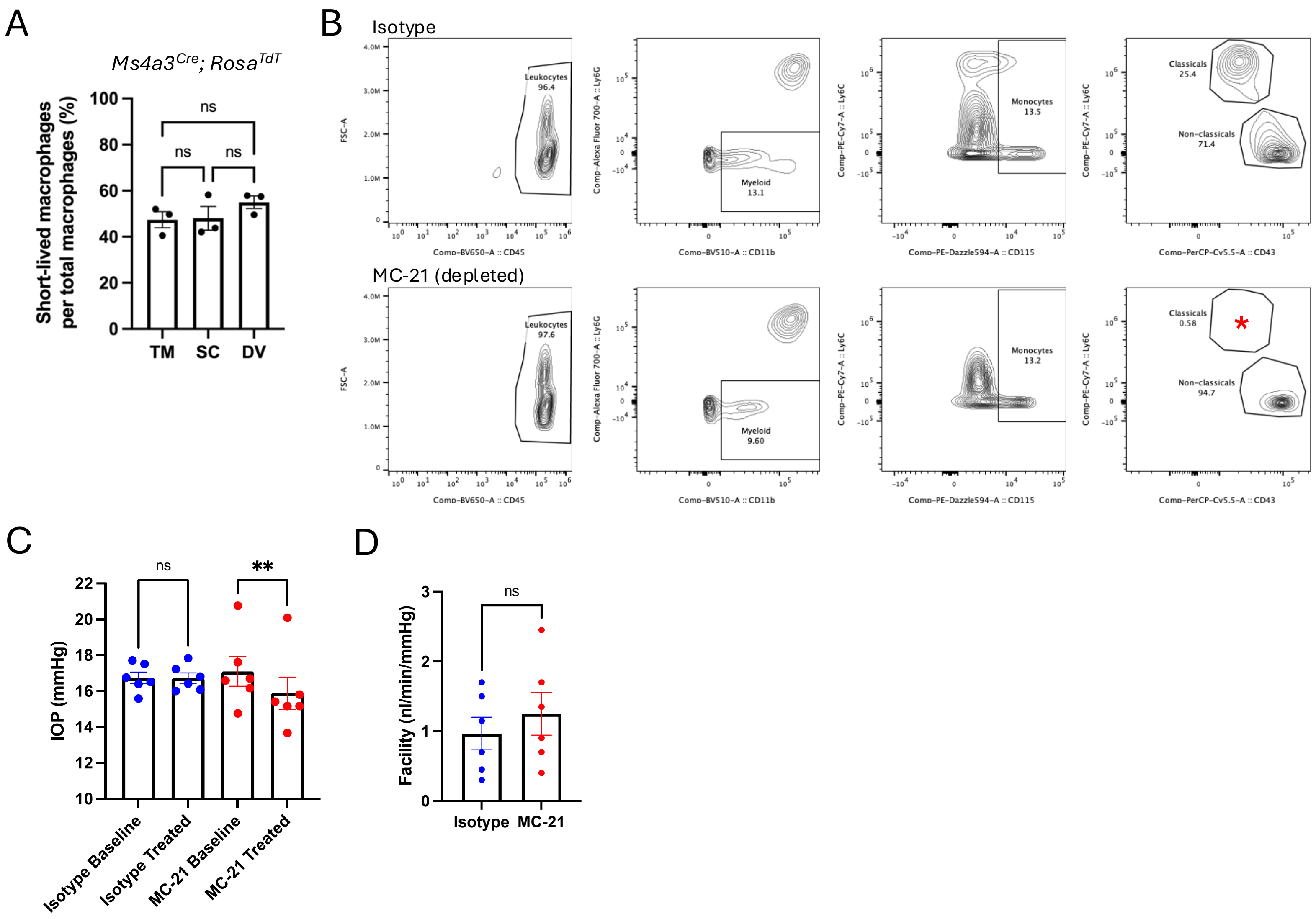

### Supplemental Figure 4

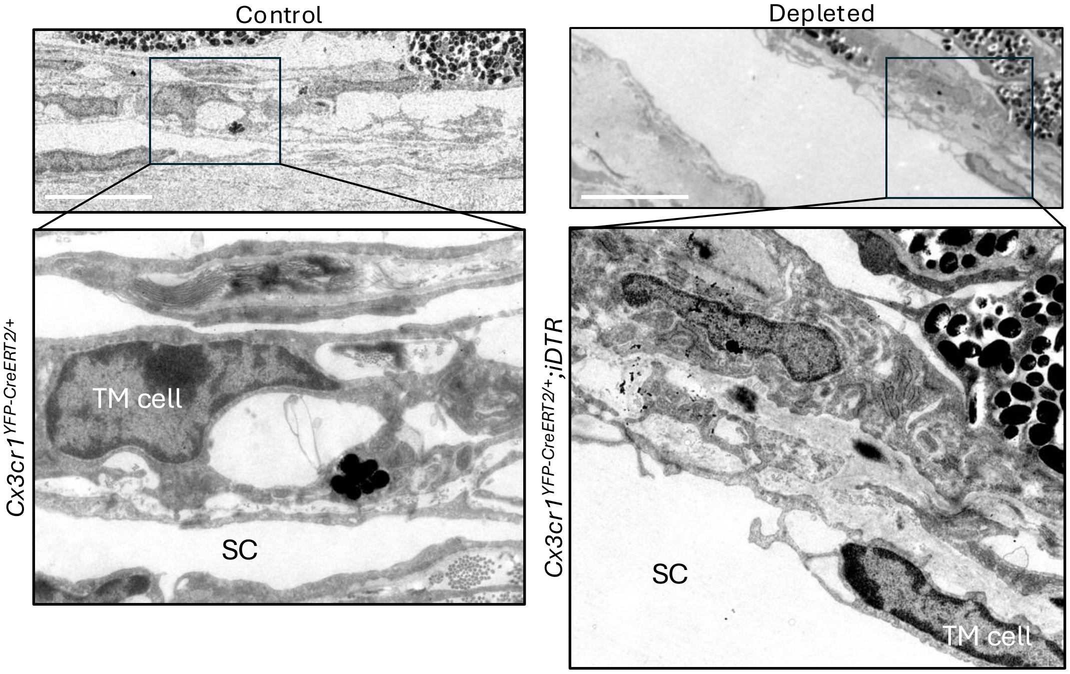

Supplemental Figure 5

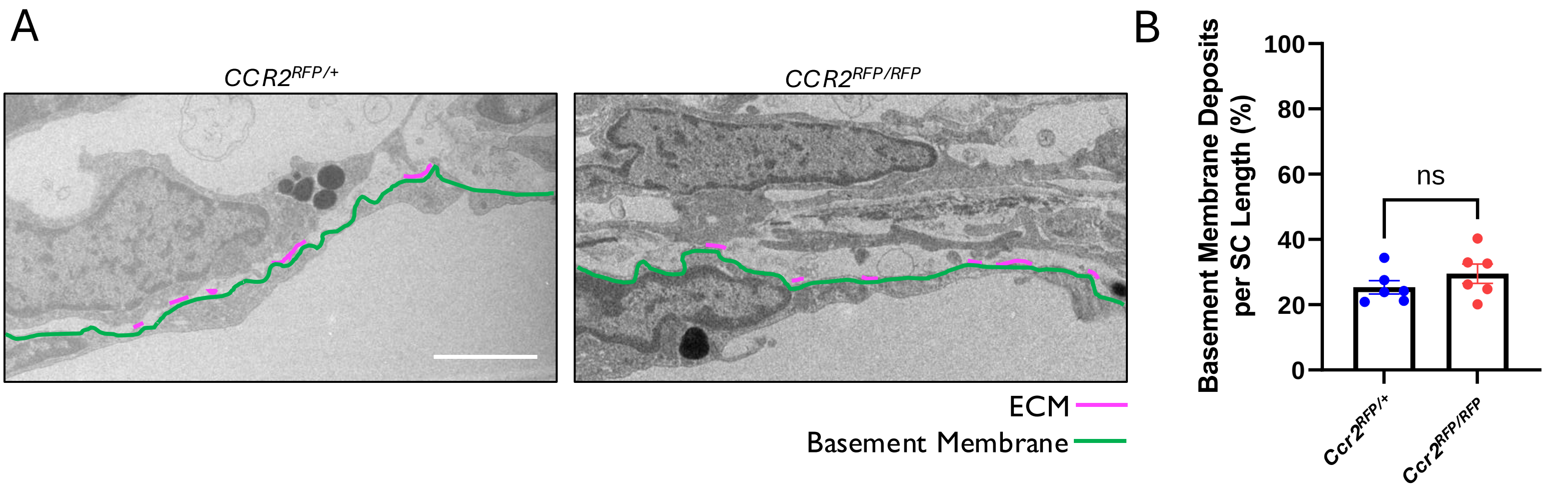
